# Mycobacterial surface-associated ESX-1 virulence factors play a role in mycobacterial adherence and invasion into lung epithelial cells

**DOI:** 10.1101/2020.10.13.337667

**Authors:** Yanqing Bao, Qi Zhang, Lin Wang, Javier Aguilera, Salvador Vazquez Reyes, Jianjun Sun

**Author notes:** Equal contribution (Yanqing Bao, Qi Zhang, Lin Wang). Corresponding author: Jianjun Sun.

## Abstract

EsxA has long been recognized as an important virulence factor of *Mycobacterium tuberculosis* (Mtb) that plays an essential role in Mtb cytosolic translocation by penetrating phagosomal membranes with its acidic pH-dependent membrane permeabilizing activity (MPA). Since the reported cytolytic activity of EsxA at neutral pH is controversial, in the present study we have obtained direct evidence that it is the residual ASB-14, a detergent used in EsxA purification, but not EsxA that causes cytolysis at neutral pH. Besides, we have also found that the exogenously added EsxA was internalized into lung epithelial cells (WI-26) and inserted into the host membranes, and these processes could be blocked by cytochalasin D and bafilomycin A. This indicates that EsxA is bound by host surface receptors and internalized into acidic endosomal compartments. This observation has intrigued us to investigate the role of EsxA in mycobacterial adherence and invasion in host cells. Interestingly, compared to the *Mycobacterium marinum* (Mm) wild type strain, the Mm strain with deletion of the *esxBA* operon (MmΔEsxA:B) had a lower adherence but a higher invasion in WI-26 cells. More interestingly, either inducible knockdown of EsxAB or removal of the bacterial surface-associated EsxAB by Tween-80 exhibited opposite results compared to gene knockout. Finally, the surface-associated EsxA is correlated to mycobacterial virulence. Together, the present study has shown for the first time that EsxA is internalized into the host cells and inserts into the host membranes, and mycobacterial surface-associated EsxAB plays an important role in mycobacterial adherence and invasion in host cells, which warrants further investigation.

## Introduction

*Mycobacterium tuberculosis* (Mtb), the causative bacterial pathogen for tuberculosis (TB) disease, infects one-third of the world’s population and causes over one million deaths worldwide each year (1,2). It is estimated that an airborne droplet carrying 1–3 Mtb bacilli is enough to cause systemic dissemination and infection (3,4). When Mtb is inhaled into the lung, it is engulfed into alveolar macrophage. Instead of being destroyed in phagolysosomes like many other intracellular pathogens, Mtb employs various virulence factors to establish intracellular survival and infection. Among them, EsxA (6 kDa early secreted antigenic target, ESAT-6) has been found necessary for Mtb virulence in several aspects, including mycobacterial cytosolic translocation, induction of immune responses and intracellular survival in host cells (5–8).

EsxA is the substrate of the ESX-1 system, a type VII secretion system that is present in Mtb but deleted in the vaccine strain BCG (9–15). EsxA is found to be secreted with its chaperon EsxB (10 kDa cell filtrate protein, CFP-10) in a co-dependent manner (16–20). Biochemical analysis has shown that EsxA and EsxB form a 1:1 heterodimeric complex (21–25). Multiple studies demonstrate that defective secretion of EsxA attenuates Mtb’s virulence, stressing the necessity of secreted EsxA (6,10,15). Moreover, emerging evidence indicate that instead of being directly released to the surrounding medium, the secreted EsxA stays associated with the mycobacterial surface, and the surface-associated EsxA plays a major role in mycobacterial virulence (26,27).

In our earlier studies with the purified EsxA and EsxB, we have characterized the pH-dependent membrane interactions and conformational changes of EsxA and EsxB and found that at acidic pH (pH 5 and below) EsxA, but not EsxB, makes a significant conformational change and permeabilizes liposomal membranes. The membrane-permeabilizing activity (MPA) is unique to EsxA from Mtb, but not present in its ortholog from non-pathogenic *Mycobacterium smegmatis* (28). The single-residue mutations at Q5 either up-regulate (e.g. Q5V and Q5L) or down-regulate (e.g. Q5K and Q5R) the MPA of EsxA *in vitro*. And these mutations accordingly either up- or down-regulate mycobacterial cytosolic translocation and virulence in macrophages and zebra fishes (29). Thus, our data strongly support that EsxA MPA contributes to mycobacterial virulence.

In addition to functioning within phagosomes to mediate mycobacterial cytosolic translocation, exogenously added EsxA protein has also been shown to cause cytolysis of red blood cells, macrophages, and type 1 and type 2 pneumocytes at neutral pH, suggesting that it may directly act on plasma membrane like a cytolytic pore-forming toxin (7,10,21,30–32). However, the data regarding to the ability of EsxA to cause cytolysis and cell death at neutral pH are controversial. In the studies assessing the effects of EsxA on immune responses, EsxA was shown to modulate the inflammatory responses of macrophages and T cells, but did not cause cytolysis or apoptosis (33–35). Refai et al reported that a zwitterionic detergent amidosulfobetaine-14 (ASB-14), which is commonly used in purification of recombinant EsxA in the protocol provided by BEI Resource, was involved in EsxA-mediated cytolysis (36). They found that the recombinant EsxA existed as a dimer or oligomer, which was not cytolytic, while the ASB-14-treated EsxA existed as a monomer with conformational changes and caused membrane-lysis and cell death (36). A recent study, however, showed that the recombinant EsxA protein didn’t lyse cell membrane and the lytic activity previously attributed to EsxA was due to the residual ASB-14 detergent in the preparation (37).

To solve the controversy, in the present study, we investigated the cytotoxic effects of ASB-14 and obtained direct evidence that it is ASB-14, not EsxA that causes cytolysis at neutral pH. More importantly, it is for the first time we found that EsxA was internalized into the cells through endocytosis, trafficked to acidic subcellular organelles and inserted into the host cell membrane. Since receptor-mediated endocytosis or phagocytosis are common pathways for bacterial pathogens to adhere and invade into host cells (38), we hypothesize that the bacterial surface-associated EsxAB may play a role in mediating mycobacterial adherence and invasion. Thus, in this present study, we have further investigated the role of EsxAB in mycobacterial adherence and invasion in epithelial cells.

## Results

### It is ASB-14, but not EsxA that causes cytolysis

To solve the controversy whether EsxA protein or ASB-14 causes cytolysis, we tested the potential cytotoxic effect of the purified EsxA protein in the presence or absence of ASB-14. The C-terminally His-tagged EsxA protein was overexpressed in the inclusion body in *E. coli*. The inclusion body was purified and solubilized by 8 M urea and then applied to a Ni-column, in which EsxA protein was refolded by slowly removing urea and eluted as a soluble protein with an imidazole gradient. No detergent was used during the purification process. We found that addition of the EsxA protein up to 40 μg/ml into the WI-26 cell culture caused little cytotoxicity (**Fig. 1A**). Next, we mixed the EsxA protein with 0.5% ASB-14 overnight and then applied the protein-detergent mixture to gel filtration chromatography using a Superdex 75 column or to dialysis using 3 kDa cut-off filter membrane. We found that the resultant EsxA protein after gel filtration or dialysis caused significant cell death even at low protein concentrations (5-10 μg/ml) (**Fig. 1A**).

**Fig 1.**
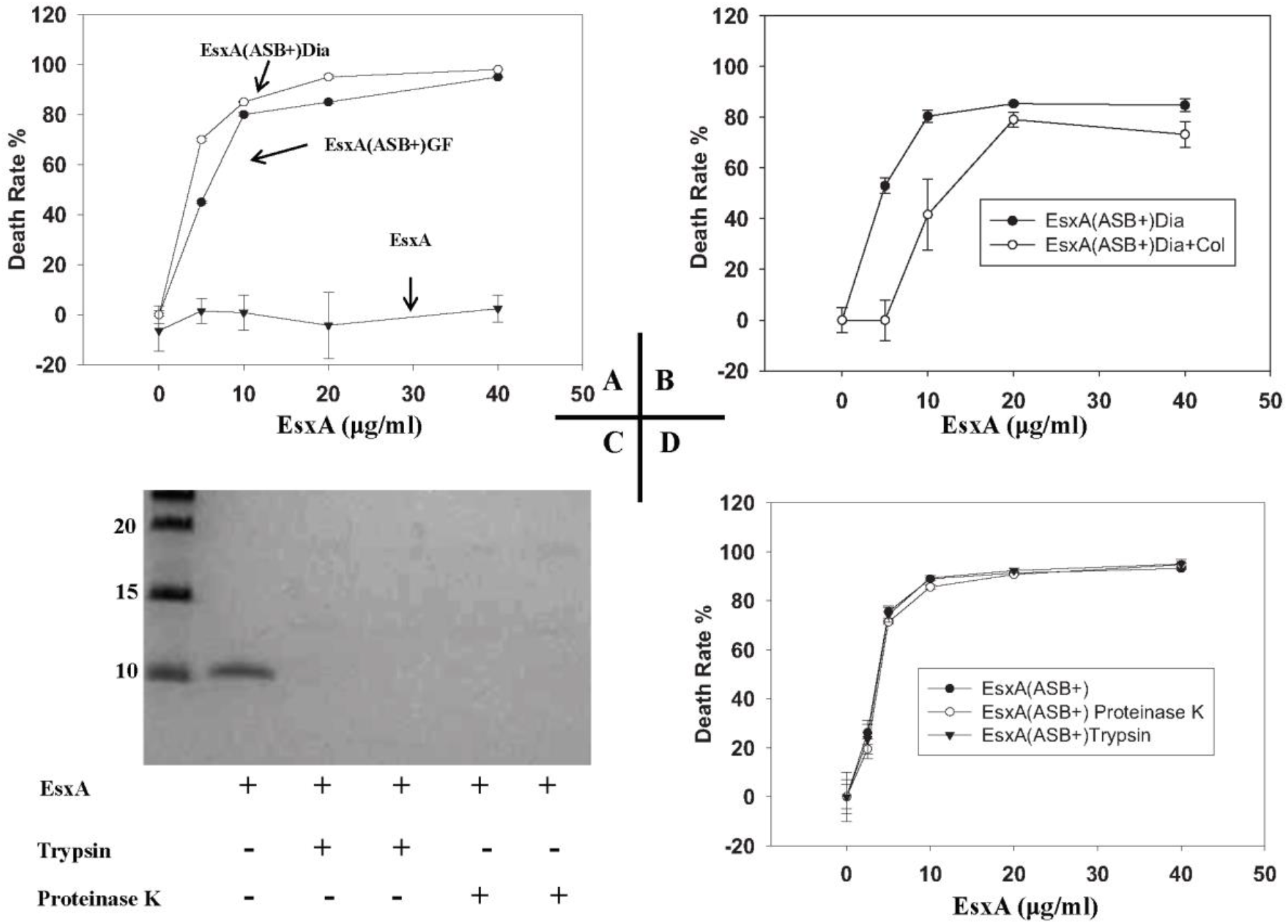
EsxA did not cause cytotoxicity in WI-26 cells. **A**. The purified EsxA protein was incubated with 0.5% ASB-14 for 4 h and then was either applied to gel filtration or dialysis to remove ASB-14. The resultant proteins were designated as EsxA(ASB+)GF or EsxA(ASB+)Dia. EsxA, EsxA(ASB+)GF and EsxA(ASB+)Dia were incubated with WI-26 cells at the indicated concentrations for 72 h. The cells that were incubated with medium alone (no protein) were set as negative controls. The cytotoxicity was measured by MTS assay and the death rate (%) was calculated as: (OD_492_ of control cells – OD_492_ of test cells)/(OD_492_ of test cells) × 100. The data were calculated from at least three independent experiments. **B**. The EsxA(ASB+)Dia sample was further passed through a HiPPR detergent removal column and the flow-through sample was collected and termed EsxA(ASB+)Dia+Col. EsxA(ASB+)Dia and EsxA(ASB+)Dia+Col were incubated with WI-26 cells for 72 h at the indicated concentrations. The cytotoxicity was measured as described above. **C**. The ASB-14-treated EsxA protein or EsxA(ASB+), was incubated with either 0.025% trypsin or 0.05% proteinase K for 30 min. The samples were applied to SDS-PAGE and stained by Coomarssie blue to confirm the digestion of EsxA protein after treatment of trypsin and proteinase K. **D.** Then the samples were incubated with WI-26 cells for 72 h. The cytotoxicity was measured as described above.

We suspected that ASB-14 was not completed removed by gel filtration or dialysis. Therefore, we applied the dialyzed EsxA protein sample to a HiPPR detergent removal column. Interestingly, while the sample treated with the HiPPR detergent removal column still caused cell death but had a significantly higher LD_50_ (~ 13 μg/ml), compared to the sample without passing through the HiPPR column (LD_50_ ~ 5 μg/ml) (**Fig. 1B**). This indicates that the ASB-14 left in the EsxA protein solution may be responsible for cell death.

To further test our hypothesis, the EsxA protein solution containing 0.5% of ABS-14 was treated with trypsin or proteinase K. While SDS-PAGE showed that the EsxA protein was completely digested (**Fig. 1C**), the digested protein solutions were still able to cause significant cell death (**Fig. 1D**). Trypsin or proteinase K alone at the applied concentration did not cause any cell death (data not shown). These data strongly suggest that it is ASB-14, but not EsxA that causes cell death, and ASB-14 can’t be efficiently removed by gel filtration, dialysis and even the detergent removal column.

### The residual contamination of ASB-14 in EsxA protein preparation is enough to cause cell lysis

To further confirm that ASB-14 is cytotoxic to the cells, we tested the dose-dependent cytotoxicity of ASB-14. As low as 5 μg/ml, ASB-14 caused over 60% cell death, and at 50 μg/ml ASB-14 caused 100% cell death (**Fig. 2A**). Now the question remains: after gel filtration, dialysis or passing through the HiPPR column, is the ASB-14 left in the protein solution enough to kill cells? We went ahead to determine the amount of ASB-14 in the EsxA protein solutions by quantitative HPLC. We first established a standard curve using the pure ASB-14 at various concentrations (**Fig. 2B**) and then measured the abundance of ASB-14 in the EsxA protein solutions (**Fig. 2C**). The identity of ASB-14 in protein solutions was confirmed by mass spectrometry. Based on the standard curve, the concentrations of ASB-14 in the protein solutions were calculated (**Table 1**). We further calculated the actual concentrations of ASB-14 that were applied into the tissue culture when the EsxA protein was used at 20 μg/ml (**Table 1**), a condition that killed at least 80% of cells (**Fig. 1A, B** **and** **D**). The actual concentrations of ASB-14 applied in the cell culture ranged from 12-32 μg/ml (**Table 1**), which were enough to cause significant cytotoxicity as shown in **Fig. 2A**. Together, the data confirme that it is ASB-14, but not EsxA that causes cell death.

**Fig 2.**
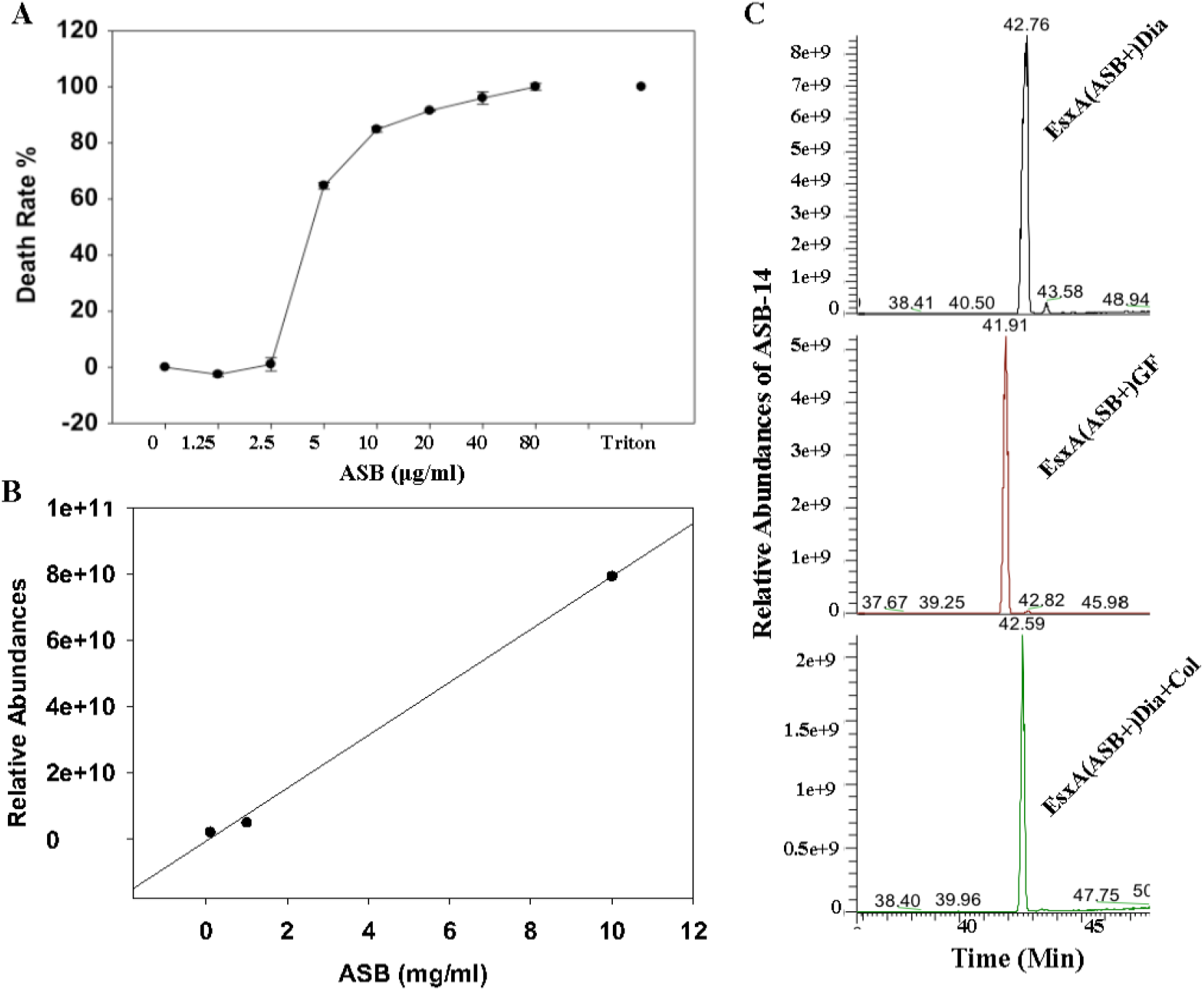
After gel filtration, dialysis, or passing through detergent removal column, the EsxA protein solution still has residual ASB-14 that is enough to kill cells. **A**. WI-26 cells were incubated with the standard solutions of ASB-14 at the indicated concentrations for 72 h. 0.1% Triton X-100 was used as a positive control. The death rate was measured and calculated as described above. **B**. The standard solutions of ASB-14 at various concentrations (0.001, 0.01, 0.1, 1, 10 mg/ml) were applied to the quantitative HPLC to establish the standard curve. **C**. The protein samples EsxA(ASB+)Dia, EsxA(ASB+)GF and EsxA(ASB+)Dia+Col were treated with 90% acetone at cold to precipitate proteins, and then the remaining acetone solutions were dried under nitrogen and the samples were applied to quantitative HPLC to quantify the abundance of ASB-14.

**Table 1.** Concentrations of ASB-14 in EsxA protein solutions measured by quantitative liquid chromatography.

| Samples | ASB-14 concentration | Final concentration of ASB-14 <sup>a</sup> |
| --- | --- | --- |
| EsxA(ASB+)GF | 1.46 mg/ml (3.36 mM) | 22.5 ug/ml (51.76 μM) |
| EsxA(ASB+)Dia | 1.93 mg/ml (4.44 mM) | 32.2 ug/ml (74.08 μM) |
| EsxA(ASB+)Dia+Col | 0.60 mg/ml (1.38 mM) | 12 ug/ml (27.6 μM) |

<sup>a</sup> The final concentrations of ASB-14 in the cultured cells, when EsxA is 20 μg/ml (2 μM).

It is worth of mentioning that we tested ASB-14 compound from three different vendors (Sigma Aldrich, Santa Cruz Biotechnology, and EMD Millipore), and all brands of ASB-14 showed similar cytotoxic effects. We also purified the EsxA protein by following exactly the protocol from BEI Resource, and we found the EsxA protein solution was cytotoxic.

### EsxA protein was internalized into the host cell through endocytosis and inserted into the membrane under an acidic condition

Since EsxA is not cytolytic, we are intrigued to investigate whether EsxA can traffic into the cells and/or insert into the host membranes. NBD, an environmental sensitive dye, has little fluorescence in aqueous solution, but emits strong fluorescence when it inserts into lipid membranes. Thus, NBD has been used as a fluorescence marker for protein membrane insertion (39–41). Earlier, we had obtained evidence that the central Helix-Turn-Helix motif of EsxA inserted into the liposomal membrane at low pH (42). For instance, when NBD was labeled at S35C, a position that was embedded in the membrane, the NBD-labeled EsxA(S35C) showed a strong fluorescence upon acidification.

However, when NBD was labeled at G10C, a position in the N-terminal arm that did not insert into the membrane, the NBD-labeled EsxA(G10C) did not emit fluorescence (42). Here, we used the NBD-labeled EsxA(S35S) and EsxA(G10C) to test the membrane insertion of EsxA within the host cells (**Fig. 3A** **and** **B**). The NBD-labeled proteins were first incubated with the cells at 4 °C, which allowed the proteins bind to the cell surface without endocytosis. Then the temperature was shifted to 37 °C, which allowed endocytosis to occur. The real-time fluorescence emission of the NBD-labeled proteins was monitored using a fluorometer. We found that upon temperature shifted from 4 °C to 37 °C, the fluorescence emission of EsxA(S35C)-NBD exhibited a significant increase over time, while EsxA(G10C)-NBD emitted little fluorescence, which is consistent to the result obtained in liposomes (42). Our recent study has identified Q5 as a key residue that regulates EsxA MPA (29). The mutation Q5K significantly reduced the MPA of EsxA and hence attenuated mycobacterial cytosolic translocation and virulence. Consistent to the result, we found that EsxA(Q5K/S35C)-NBD had a significantly lower fluorescence emission inside the cells (**Fig. 3A** **and** **B**). As expected, addition of cytochalasin D, which inhibits endocytosis by blocking actin polymerization, significantly reduced the fluorescence emission of EsxA(S35C)-NBD. Bafilomycin A, an inhibitor of endocytic acidification, exhibited a similar inhibitory effect. Together, the data suggest that EsxA is internalized into the host cell through endocytosis and inserts into the endosomal membranes in an acidic condition.

**Fig 3.**
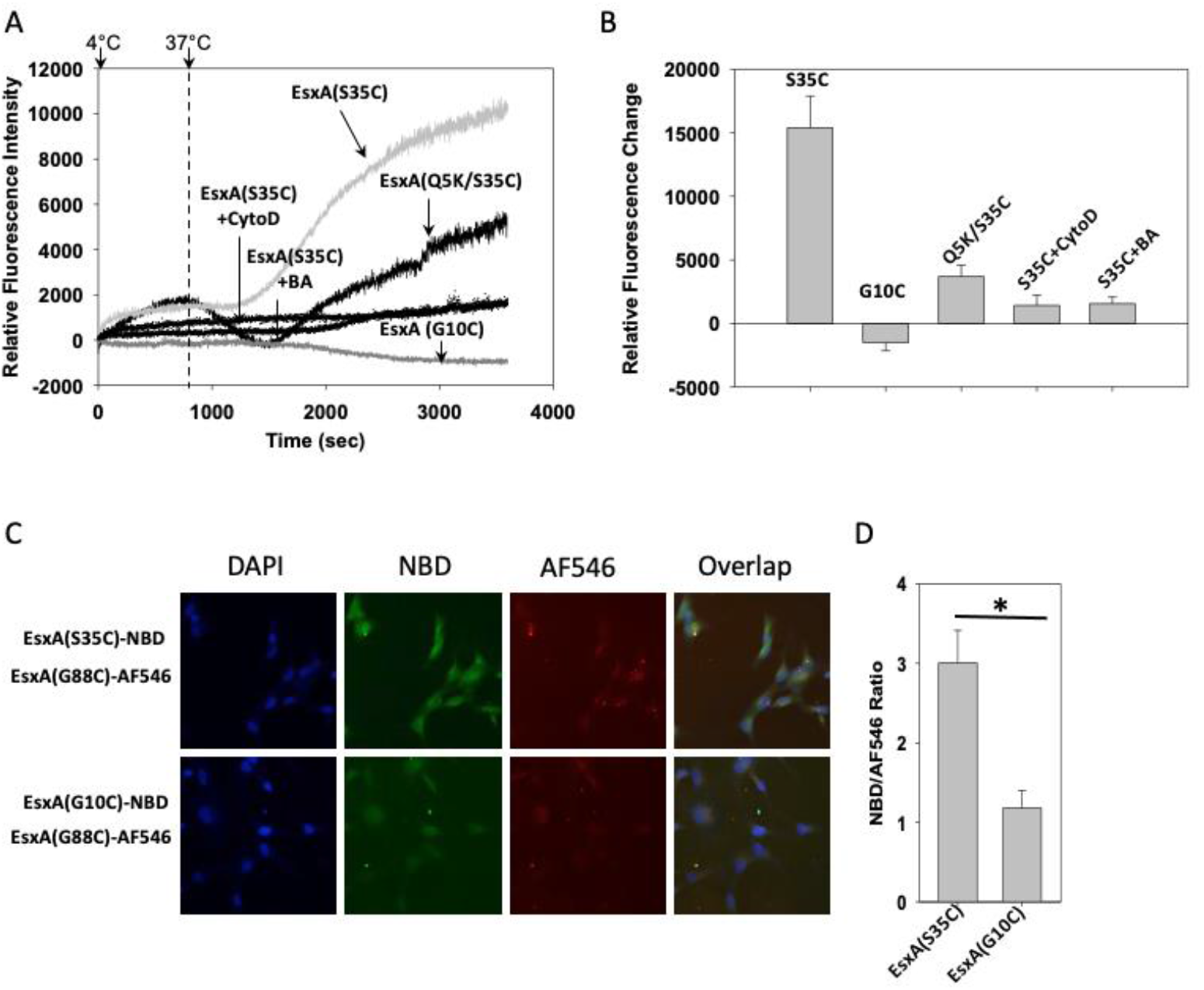
EsxA was internalized into the WI-26 cells through endocytosis and inserted into the host membrane. **A.** The suspended WI-26 cells were incubated on ice with the NBD-labeled EsxA(S35C), EsxA(Q5K/S35C) or EsxA(G10C), respectively. After 30 min of incubation, the cells were washed and suspended in cold PBS and transferred to a cuvette with magnetic stirring in the cuvette chamber of a fluorometer. The cuvette chamber was pre-chilled at 4 °C by connecting to circulating water bath. At time zero, the temperature of the water bath was set to 37 °C, and the fluorescence intensity of NBD (Ex at 488 nm, Em at 545 nm) was recorded with time. To inhibit endocytosis, WI-26 cells were pre-treated with 50 μM cytochalasin D (CytoD) or 1 μM bafilomycin A1 (BA) for 35 min at 37 °C. The cells were then incubated with the NBD-labeled EsxA protein as described above. The representative NBD emission curves with time are shown. **B.** The relative increase of NBD fluorescence intensity at 1 h (3600 seconds) incubation was shown. The data were calculated from three independent experiments. **C.** The WI-26 cells were incubated on ice with either a 1:1 mixture of EsxA(S35C)-NBD and EsxA(G88C)-AF546 or a 1:1 mixture of EsxA(G10C)-NBD and EsxA(G88C)-AF546 for 1 h. The cells were washed and then transferred to a CO_2_ incubator at 37 °C for 1 h. The cells were then fixed with 4% PFA and stained with DAPI. Finally, the cells were imaged under a fluorescence confocal microscope at three separate channels, blue (DAPI), green (NBD) and red (AF546). **D**. The intensities of NBD emission from EsxA(S35C) and EsxA(G10C) were measured and averaged from at least 6 random fields and the NBD intensity was normalized using the intensity of EsxA(G88C)-AF546 as an internal reference. The data was obtained from three independent experiments and were presented as mean ± S.E. (n = 3, *p < 0.05).

To directly visualize membrane insertion of EsxA inside the cells, we monitored the intracellular trafficking of EsxA(S35C)-NBD and EsxA(G10C)-NBD by confocal fluorescence microscopy (**Fig. 3C** **and** **D**). NBD emits green fluorescence when it inserts into the lipid membranes. To quantify the relative NBD fluorescence intensity of EsxA(S35C)-NBD and EsxA(G10C)-NBD, we used the AF546-labeled EsxA(G88C), which constitutively emits red fluorescence, as an internal reference of fluorescence intensity. Equal amount of EsxA(G88C)-AF546 was pre-mixed with EsxA(S35C)-NBD and EsxA(G10C)-NBD, respectively, followed by incubation with the cells. Thus, EsxA(G88C)-AF546 was co-trafficked with the NBD-labeled proteins, and its intensity was used to normalize the NBD fluorescence. Consistent to the result obtained in the fluorometer (**Fig. 3A** **and** **B**), the fluorescence intensity of EsxA(S35C)-NBD was significantly higher than that of EsxA(G10C)-NBD, which confirms the membrane insertion of EsxA within the host cells (**Fig. 3C** **and** **D**).

### Knockout of EsxAB increased the invasion efficiency on WI-26 cells

The observation that the exogenously added EsxA was internalized into WI-26 and inserted into the host membrane suggests that EsxA may bind to specific cell surface receptors and enter WI-26 cells through endocytosis pathway. This has intrigued us to investigate if EsxAB plays a role in mycobacterial adherence and invasion into host cells. Compared to Mm(WT) strain and the complemental strain MmΔEsxA:B(WT), the MmΔEsxA:B strain had a decreased adherence (**Fig. 4A**), indicating that EsxAB plays a role in assisting mycobacterial adherence to host cells. Interestingly, however, MmΔEsxA:B exhibited a significantly higher invasion than Mm(WT) and MmΔEsxA:B(WT) (**Fig. 4B**), which suggests that EsxAB may play a role in resistance to the uptake by host cells. To calibrate out the adherence background, we calculated the ratio of invasion/adherence and the results showed that MmΔEsxA:B has the highest invasion/adherence ratio (**Fig. 4C**). Therefore, it seems that EsxAB facilitates mycobacterial adherence to host cells, but inhibits the uptake by host cells. This is an exciting new finding regarding to the role of EsxAB, which has not been reported before.

**Fig 4.**
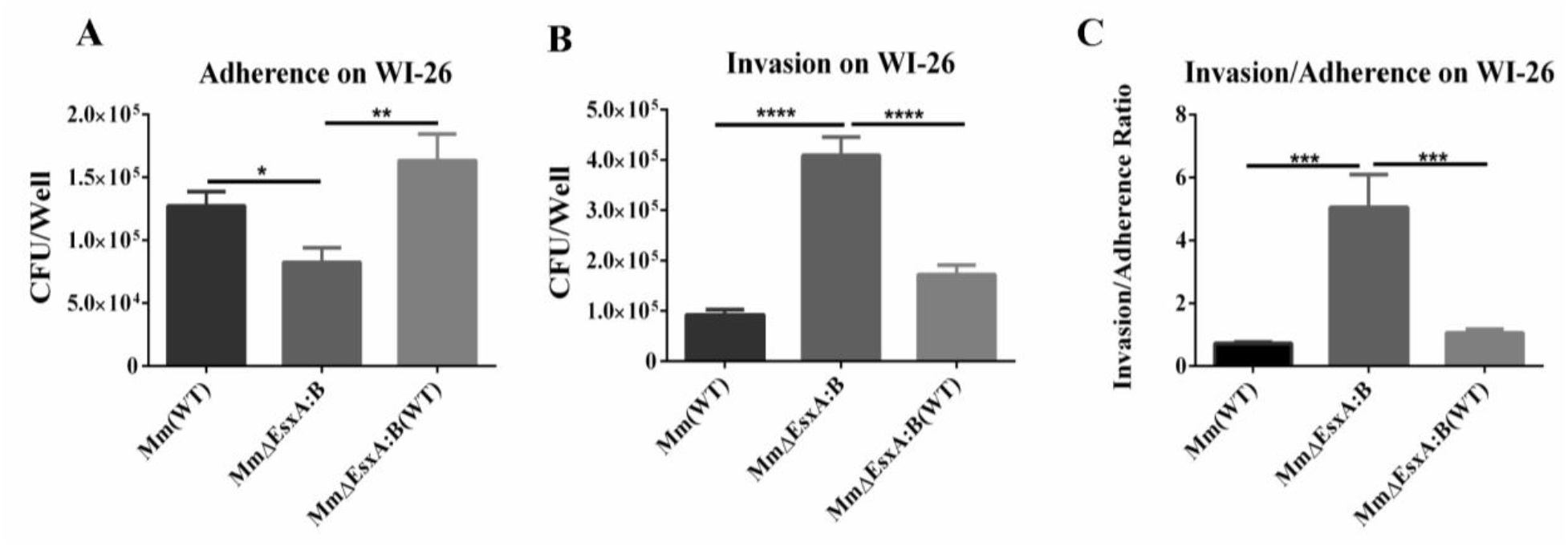
Effects of EsxAB knockout on Mm adherence and invasion into WI-26 cells. The indicated Mm strains were all cultured in 7H9 media containing 0.05% Tween-80. **A.** The WI-26 cells were pre-fixed with 4% PFA and then incubated with Mm(WT), MmΔEsxA:B and MmΔEsxA:B(WT) at MOI = 1 at 30 °C for 45 min. The cells were washed with PBS to remove non-adherent bacteria. Then the cells were harvested with 0.01% Triton X-100, and spread to 7H10 plates for CFU counting. **B**. The WI-26 cells were infected with Mm(WT), MmΔEsxA:B and MmΔEsxA:B(WT) at MOI = 1 at 30 °C for 45 min. The cells were washed by PBS to remove unbound bacteria and then incubated in DMEM medium containing 100 μg/ml Amikacin for 2 h to kill extracellular bacteria. The cells were harvested with 0.01% Triton X-100, and spread to 7H10 plates for CFU counting. **C.** The ratios of Invasion/Adherence of Mm(WT), MmΔEsxA:B and MmΔEsxA:B(WT) in WI-26 cells were calculated from the data of **Fig 4A** and **4B**. The experiment was replicated three times, in each replication there were three replicate wells for each strain. The data is presented as mean ± SD. The statistical analysis was performed with One-way ANOVA, followed by Holm-Sidak multiple comparison. \**P*<0.05, \*\**P*<0.01, \*\*\**P*<0.001, \*\*\*\**P*<0.0001.

### ATC-induced knockdown of EsxB-DAS4+ attenuated Mm’s invasion efficiency on WI-26 cells

Previous studies have shown that other factors (e.g. EspA, EspB, EspC) are co-dependently secreted with EsxAB (20,43). Thus, deletion of EsxAB may also affect the secretion of these factors. In order to determine the exact role of EsxAB in adherence and invasion without apparently affecting other factors, we used the ATC inducible protein knockdown system that has been established in the lab and reported before (44). This system allows inducible knockdown of EsxAB at the post-translational level. The Mm(EsxB-DAS4+)|pGMCKq1 cells were pre-treated with ATC (1.5 μg/ml) and then applied to infection of WI-26 cells. Surprisingly, however, the ATC-treated Mm(EsxB-DAS4+)|pGMCKq1 showed a higher adherence and a lower invasion than the group without ATC treatment (**Fig. 5A** **and** **B**). Thus, the invasion efficiency (invasion/adherence) of the ATC-treated Mm(EsxB-DAS4+)|pGMCKq1 was significant lower than that of the group without ATC treatment (**Fig. 5C**). As a control group, the adherence, invasion and invasion efficiency of Mm(WT) were all not affected by ATC treatment (**Fig. 5A-C**). Taken together, the ATC-induced knockdown of EsxAB increases adherence, but decreases invasion efficiency, which is opposite to the phenotype of EsxAB knockout.

**Fig 5.**
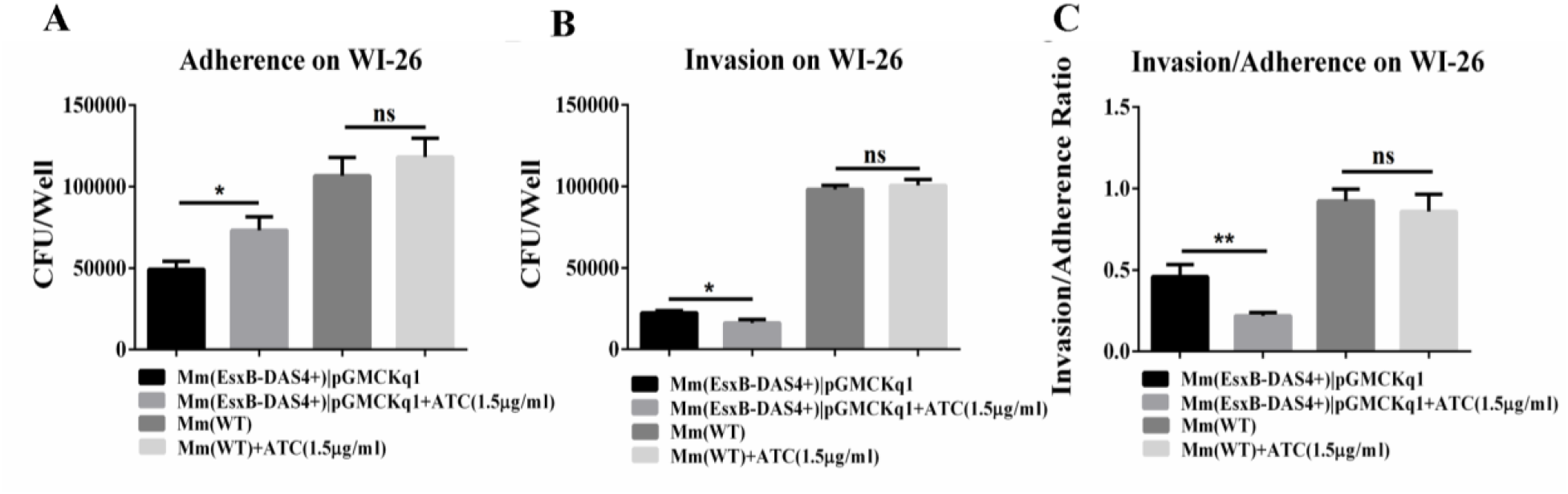
Effects of inducible knockdown of EsxAB on Mm adherence and invasion in WI-26 cells. The indicated Mm strains were cultured in 7H9 media containing 0.05% Tween-80. **A**. The WI-26 cells were pre-fixed with 4% PFA and then incubated with Mm(EsxB-DAS4+)|pGMCKq1, ATC-treated Mm(EsxB-DAS4+)|pGMCKq1, Mm(WT) and ATC-treated Mm(WT) at MOI = 1 at 30 °C for 45 min. The cells were washed with PBS to remove non-adherent bacteria. Then the cells were harvested with 0.01% Triton X-100, and spread to 7H10 plates for CFU counting. **B**. The WI-26 cells were infected with Mm(EsxB-DAS4+)|pGMCKq1, ATC-treated Mm(EsxB-DAS4+)|pGMCKq1, Mm(WT) and ATC-treated Mm(WT) at MOI = 1 at 30 °C for 45 min. The cells were washed by PBS to remove unbound bacteria and then incubated in DMEM medium containing 100 μg/ml Amikacin for 2 h to kill extracellular bacteria. The cells were harvested with 0.01% Triton X-100, and spread to 7H10 plates for CFU counting. **C**. The ratio of Invasion/Adherence was calculated from the data in **Fig. 5A** and **5B**. The experiment was replicated three times, in each replication there were three replicate wells for each strain. The data is presented as mean ± SD. The statistical analysis was performed with multiple *t*-test. \**P*<0.05, \*\**P*<0.01.

### Addition of Tween-80 reduced mycobacterial-associated EsxA

To further confirm if reduction of EsxAB could affect mycobacterial adherence and invasion, Tween-80 was used to reduce mycobacterial surface-associated EsxA (26,45). In order to efficiently and specifically label and quantify the surface associated EsxA, we used the SpyTag (ST)-SpyCatcher (SC) system that has been described in our previous report (44). Briefly, the 13-amino acid SpyTag is engineered into the C-terminus of EsxA, which produces an EsxA-ST fusion protein that has normal expression, secretion and membrane permeabilization activity as EsxA wild type. The secreted EsxA-ST can react with SC-GFP to form a fusion protein EsxA-ST-SC-GFP through a covalent bond between ST and SC (46). This covalent modification is highly specific and stable, thus it is an efficient way of labeling and quantitating the surface-associated EsxA. First, the live Mm(EsxA-ST) cells were incubated with SC-GFP to label the mycobacterial surface-associated EsxA-ST. After washes, the intensity of GFP fluorescence on the mycobacterial cells was quantified, which is directly related to the amount of the surface-associated EsxA-ST. The result showed that in the presence of Tween-80, the GFP intensity on the mycobacterial cells was much lower than those cells in the absence of Tween-80, suggesting that Tween-80 treatment has reduced the amount of the surface-associated EsxA-ST (**Fig. 6A** **and** **B**). Quantitative result showed that the green/red overlap rate of the mycobacterial cells without Tween-80 was ~ 5-fold of that with Tween-80. Similar result was acquired with by Western blots using anti-GFP antibody (**Fig. 6C** **and** **D**). At all of the tested concentrations of SC-GFP, the Mm(EsxA-ST) cells without Tween-80 showed more SC-GFP labeled EsxA-ST than that with Tween-80 (**Fig. 6C**). Calculation of the relative band intensity (EsxA-ST-SC-GFP vs GroEL) indicated that addition of Tween-80 in 7H9 media reduced ~30% of surface-associated EsxA-ST (**Fig. 6D**). The calculation data was showed in **Table 2**.

**Fig 6.**
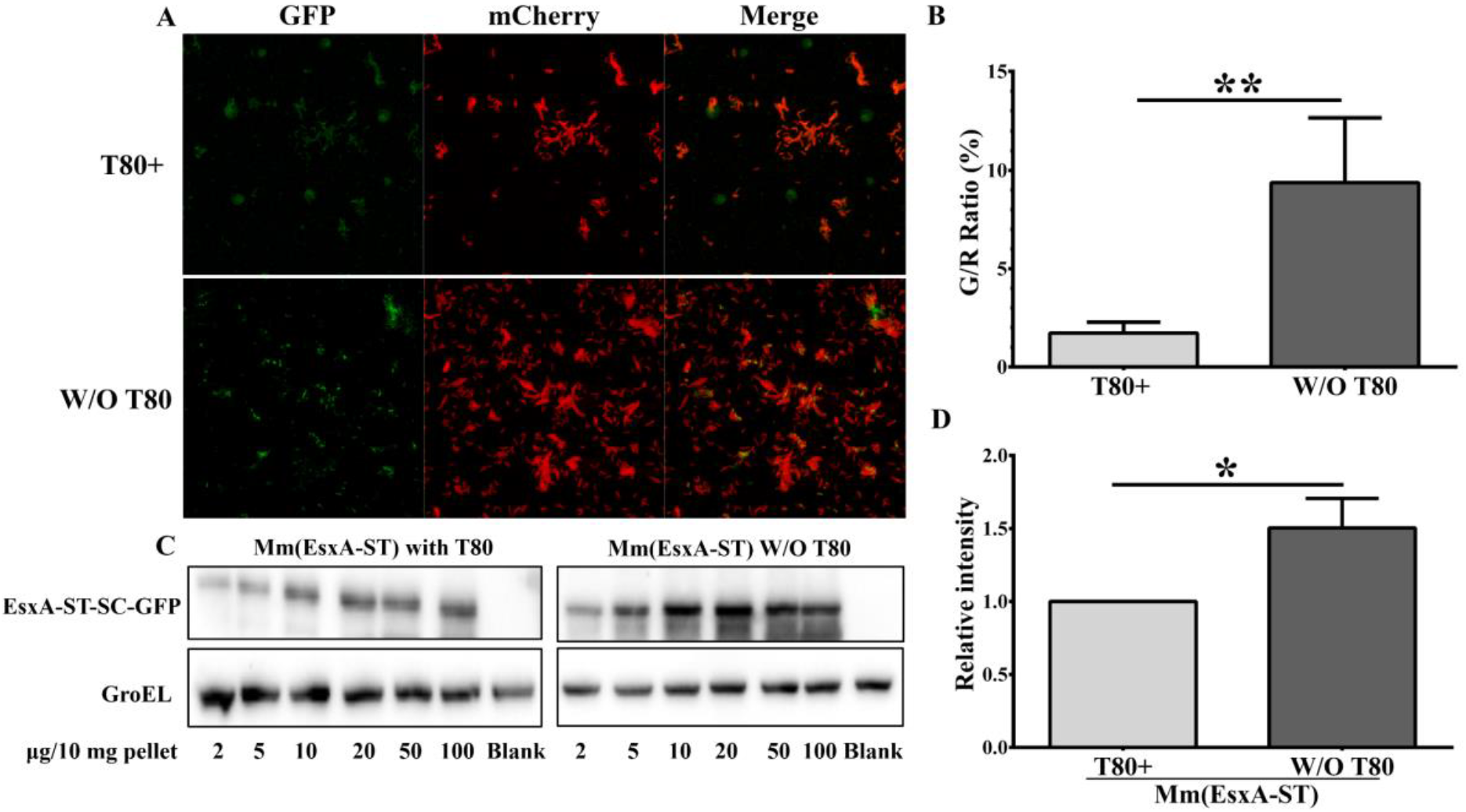
Tween-80 reduced the surface-associated EsxA. **A**. The Mm(EsxA-ST) strain expressing mCherry was incubated with SC-GFP in the 7H9 media in the presence or absence of 0.05% Tween-80. The bacteria were washed with PBS to remove free SC-GFP and subjected to confocal fluorescence microscope. The images were taken with green and red fluorescence channels. **B**. The Green/Red ratio was calculated by dividing the total green intensity with red intensity in random fields. **C**. The Mm(EsxA-ST) cells were incubated with the purified SC-GFP to label the mycobacteria-associated EsxA-ST. The Mm(EsxA-ST) was cultured in 7H9 media in the presence or absence of Tween-80. GroEL was used as a loading control. The max relatively intensity of EsxA-ST-SC-GFP in each group was used for following comparison. **D**. Relative level of mycobacterial-associated EsxA-ST in Mm(EsxA-ST). Addition of Tween-80 reduced the amount of the mycobacterial-associated EsxA-ST by ~ 30%. For fluorescence microscopy, 10 random sights were taken for each strain, and the assay was replicated for three times. The WB assay was replicated three times. The data is presented as mean ± SD. Statistical analysis was performed with *t*-test. \**P*<0.05, \*\**P*<0.01.

**Table 2.** Summary of bands’ relative intensity from WB ^a^.

| Media with/without Tween-80 | Relative intensity of each SC-GFP concentration <sup>b</sup> |  |  |  |  |  |
| --- | --- | --- | --- | --- | --- | --- |
|  | 2 | 5 | 10 | 20 | 50 | 100 |
| With Tween-80 | 1.0 | 1.298 | 2.325 | 2.777 | 2.584 | 2.867 |
|  | 1.0 | 0.697 | 2.843 | 3.941 | 5.086 | 4.659 |
|  | 1.0 | 3.039 | 2.440 | 8.016 | 8.908 | 8.293 |
| Without Tween-80 | 2 | 5 | 10 | 20 | 50 | 100 |
|  | 1.0 | 1.765 | 2.466 | 2.504 | 1.512 | 1.186 |
|  | 1.0 | 1.419 | 1.671 | 1.414 | 1.129 | 0.685 |
|  | 1.0 | 2.125 | 7.770 | 12.721 | 7.056 | 3.972 |

<sup>a</sup> The results of three replicates are included, the intensity at 2 µg/10 mg was used as control to calculate relative intensity in each replicate. The highest relative intensity in each group was used to calculate the difference of mycobacteria-associated EsxA upon treatment of Tween-80.
<sup>b</sup> SC-GFP was added into Mm(EsxA-ST) at concentrations of µg per 10 mg of bacterial pellet.

### Addition of Tween-80 attenuated mycobacterial invasion and cytotoxicity on WI-26 cells

The single-cell preparation of Mm(EsxA-ST) cultured with/without Tween-80 was used for adherence and invasion assays on WI-26 cells. The results showed that addition of Tween-80 had no effect on Mm(EsxA-ST)’s adherence, but had a significant effect on its invasion (**Fig. 7A**). While for MmΔEsxA:B, both adherence and invasion were not affected by Tween-80 (**Fig. 7B**), suggesting that the decreased invasion by Tween-80 was specific to EsxAB.

**Fig 7.**
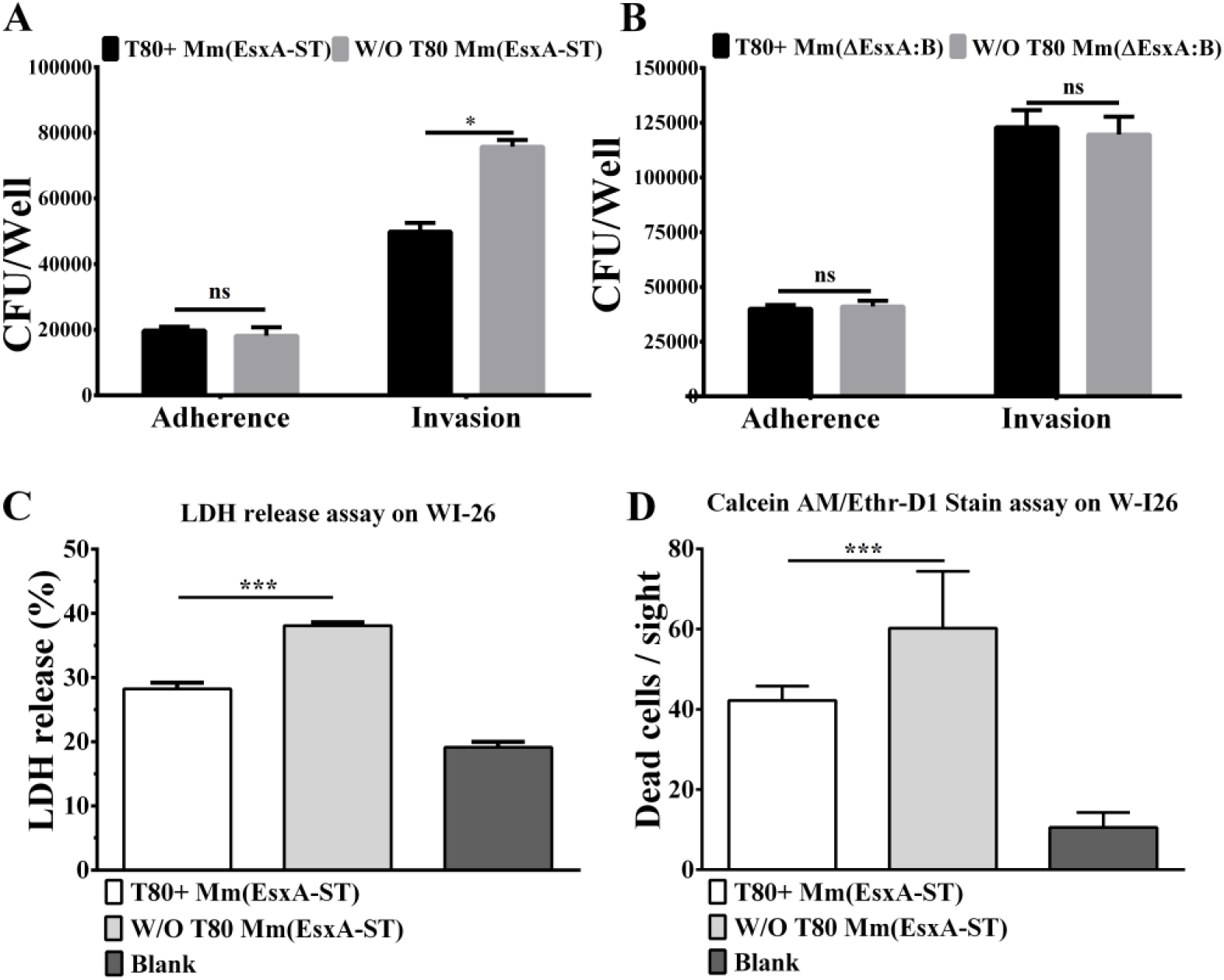
Effect of Tween-80 on adherence, invasion and cytotoxicity of Mm(EsxA-ST) and MmΔEsxA:B. The strains were cultured in the 7H9 media with/without 0.05%Tween-80, respectively. **A**. Adherence and invasion of Mm(EsxA-ST) on WI-26 cells. **B**. Adherence and invasion of MmΔEsxA:B on WI-26 cells. The experiment was replicated three times, in each replication there were three technical replicate wells for each strain. The data is presented as mean ± SD. The statistical analysis was performed with multiple *t*-test. \**P*<0.05. **C**. Cytotoxicity of Mm(EsxA-ST) cultured with/without 0.05% Tween-80. The assay was conducted with Calcein AM/Ethr-D1 method. **D.** Cytotoxicity of Mm(EsxA-ST) cultured with/without Tween-80. The assay was conduct with LDH release method. Mm(WT) cultured without Tween-80 was used as positive control. The cells without infection were used as blank control. For LDH release assay, the experiment was replicated three times, in each replication there were three technical replicate wells for each strain. For Calcein AM/Ethr-D1 stain assay, 18 random sights were taken for each strain and the assay was replicated for three times. The data is presented as mean ± SD. The statistical analysis was performed with One-way ANOVA, followed by Holm-Sidak multiple comparison. \*\*\**P*< 0.001.

As we previously reported, the MPA of EsxA is directly related to mycobacterial cytotoxicity. Down-regulation of the MPA lead to an attenuated cytotoxicity (29). So we supposed that since Tween-80 reduces the surface-associated EsxA, it would also attenuate mycobacterial cytotoxicity. Here, we used two methods to test Mm(EsxA-ST)’s cytotoxicity on WI-26 cells. Both Calcein AM/Ethr-D1 stain assay and LDH release assay showed that Tween-80 addition decreased Mm(EsxA-ST)’s cytotoxicity on WI-26 (**Fig. 7C** **and** **D**).

## Discussion

In our earlier biochemical characterization, we have found that EsxA requires acidic pH (≤ 5) to permeabilize liposomal membranes (28,42). Recently, we have characterized the single-residue mutations at Q5 of EsxA (e.g. Q5K and Q5V) and demonstrated that the acidic pH-dependent MPA of EsxA is required for mycobacterial cytosolic translocation and virulence (29). Most recently, we have found that the N^α^-acetylation of EsxA at T2 is essential for the low pH-dependent EsxAB heterodimer dissociation, which is the prerequisite for EsxA membrane insertion. Single-residue mutations at T2 (e.g. T2A and T2R) disrupted N^α^-acetylation, resulting in attenuated cytosolic translocation and virulence (47). All these findings support that EsxA MPA functions within the acidic subcellular compartments and contributes to the virulence of Mtb through rupturing the host phagosome membranes. Interestingly, several studies have reported that the exogenously added recombinant EsxA acted directly on plasma membrane and caused cytolysis at neutral pH, suggesting that EsxA could function as a cytolytic pore-forming toxin (7,10,21,30–32). On the contrary, however, several other studies didn’t observe any cytolytic effect of the exogenously added EsxA on the cultured mammalian cells (33–35).

In the present study, we further investigated this discrepancy and obtained evidence that ASB-14, the detergent used in EsxA preparation, is responsible for the observed cytolysis. ASB-14 is a zwitterionic detergent and useful for extraction of membrane proteins. ASB-14 has molecular weight 434.68 and critical micelle concentration at 8 mM. When 0.5% (11.5 mM) of ASB-14 was used in EsxA purification, even after gel filtration, dialysis and passage through the HiRPP detergent removal column, there was still significant amount of ASB-14 (1.38-4.44 mM) present in the protein solution (**Table 1**). By comparing the ASB-14 cytotoxicity titration curve, the residual ASB-14 was enough to cause significant cytolysis (**Fig. 2**). Another evidence comes from the digestion experiment. Even though EsxA was completely digested by the proteases, the solution containing ASB-14 was still cytotoxic (**Fig. 1C** **and** **D**). Therefore, this study has further clarified the controversy and demonstrated that EsxA doesn’t cause cytolysis on plasma membrane at neutral pH. This is consistent with our earlier studies that EsxA requires acidic pH to permeabilize the membrane (28).

Instead of causing cytolysis, we found that EsxA was internalized into the host cell through endocytosis and trafficked to the acidic subcellular organelles, where it inserts into the membranes (**Fig. 3**). To the best of our knowledge, it is for the first time that EsxA acidic pH-dependent membrane insertion is observed inside the host cells, which is consistent to the results obtained in liposomes (28,29,42,48). Emerging evidence have demonstrated that EsxAB stays on the surface of mycobacteria and contributes to the virulence (26,27,49,50). Besides, on epithelial cells, endocytosis is usually related to bacteria’s invasion (21,22). Thus, this result has intrigued us to investigate if the surface-associated EsxAB mediates mycobacterial adherence and invasion in host cells.

As a virulence factor, lack of EsxA results in attenuation of mycobacterial virulence (12,29,51). Interestingly, compared to Mm(WT), MmΔEsxA:B had a lower adherence and a higher invasion (**Fig. 4**). This indicates that EsxAB may play a positive role in mycobacterial adherence, but a negative role in invasion. Surprisingly, however, inducible knockdown of EsxAB by the DAS4+ system showed the exactly opposite results as knockout of EsxAB (**Fig. 5**). Moreover, reduction of the surface-associated EsxAB by Tween-80 had similar results as the inducible knockdown (**Fig. 6** **and** **7**). Thus, the data suggest that in the absence of the *esxBA* operon (e.g. MmΔEsxA:B), mycobacteria use an EsxAB-independent mechanism for invasion, while in the presence of the operon (e.g. inducible knockdown and Tween-80), mycobacteria use an EsxAB-dependent mechanism for invasion. While it is reasonable to believe that there may be a compensatory mechanism in the MmΔEsxA:B strain, the result showed that the complemental strain MmΔEsxA:B(WT) restored the wild type phenotypes seems to argue against the presence of a genetic compensatory mechanism (**Fig. 4**). There are several mycobacterial factors whose expression and/or secretion are dependent on EsxAB. Thus, deletion of the *esxBA* operon has a potential impact on expression or secretion of these factors (20,43). The roles of these factors in mycobacterial adherence and invasion warrant further investigation.

Despite of the opposite results, the experiments with either knockout or knockdown of EsxAB strongly suggest that EsxAB play an important role in mycobacterial adherence and invasion, presumably through the bacterial surface-associated EsxAB that binds to the surface receptors of the host cells and mediates bacterial invasion via receptor-mediated phagocytosis. While the surface receptors for EsxAB have not been identified yet, several earlier studies have presented evidence that EsxAB interacted with various host proteins. Earlier, Renshaw et al. found that the fluorescently labeled EsxAB heterodimer bound to the surfaces of monocytes and macrophages, but not fibroblasts (25). Later, EsxA was shown to directly bind to TLR2, which inhibited TLR signaling in macrophages (35). In a separate study, however, EsxA was shown to bind to type 1 and type 2 pneumocytes and purified human laminin (31). Recently, in a yeast two-hybrid screening, EsxA was found to bind directly to β2-microglobulin (β2M) and enter ER where it sequestered β2M to downregulate class-I-mediated antigen presentation (52). Of note, according to our data on other cell lines, deletion of EsxAB has the similar result in THP-1 cell line as in WI-26, but it has no influence on invasion efficiency in RAW264.7 and A549 cell lines (**Fig S1**), which indicates the role of EsxAB in adherence and invasion has cell-type specificity. In future, more in-depth studies are needed to investigate the mechanism of EsxA-mediated pathogen-host interaction, such as identification of the receptors and intracellular trafficking pathways.

## Experimental procedures

### Bacterial strains and cell lines

The Mm(WT) strain was purchased from American Type Culture Collection (ATCC, Manassas, VA, USA) and preserved in our lab. The Mm strain with deletion of *esxBA* operon (MmΔEsxA:B) and the complemental strain (MmΔEsxA:B(WT)) were preserved in our lab (53). The WI-26 VA4 cell line, RAW264.7 cell line and THP-1 cell line were purchased from ATCC and preserved in our lab. The A549 cell line was generously offered by Dr. Jianying Zhang from University of Texas at El Paso. The strains of Mm(EsxA-ST) and Mm(EsxB-DAS4+)|pGMCKq1 were generated and preserved in our lab as described before (44).

### Preparation of single-cell bacterial culture

The single-cell bacterial culture for infection assay was prepared as described elsewhere (54). Briefly, the Mm strains were inoculated into 100 ml 7H9 (10% OADC, 0.5% glycerol, with or without 0.2% Tween-80), and cultured at 30 °C until OD_600_ reached 0.6-0.8. Then, the bacteria was collected by centrifugation and washed twice with phosphate saline buffer (PBS) and resuspended with 3 to 5 ml of the same 7H9 media used for culture. The resuspended culture was pushed through 27-gauge needle several times to break the clumsy, which followed by a passage through 5μm filter to further isolate the cells. The aliquots were stored at −80 ºC and the concentration of the single cell preparation was determined by colony forming unit (CFU) in 10-fold serial dilutions.

### Protein expression and purification

EsxA was expressed and purified from *E. coli* BL21(DE3) as described previously (28). Briefly, the inclusion body was isolated and then solubilized in 8 M urea. The proteins were refolded on a nickel column and eluted with an imidazole gradient. The eluted proteins were further purified by size exclusion chromatography using a Superdex 75 column that was pre-equilibrated with the buffer 20 mM TrisHCl, 100 mM NaCl, pH 7.3.

The EsxA mutants EsxA(S35C), EsxA(G10C) and EsxA(G88C) were generated by site-directed mutagenesis and purified as previously described (42). The EsxA(Q5K/S35C) was expressed and purified as a GST-fusion protein in BL21 (DE3), followed by cleavage off the GST tag as previously described (28).

For SC-GFP, the plasmid pQE80L-SC-ELP-GFP was purchased from Addgene (#69835, Watertown, MA, USA). The ELP linker between SC and GFP was deleted by PCR using the primers that flank SC and GFP. The amplified fragment was inserted into EcoR I and Sac I sites of the pQE80L. The His-tagged SC-GFP fusion protein was expressed in BL21(DE3) and purified in a Ni^2+^-affinity column in an AKTA FPLC as described elsewhere (GE Lifesciences, Chicago, IL, USA) (55).

### Preparation of ASB-14-treated EsxA protein

The purified EsxA protein was incubated with 0.5% ASB-14 overnight on ice. Then the samples were either applied to size exclusion chromatography using a Superdex 75 column or dialyzed in a Spectra/Por3 dialysis bag with 3 kDa MWCO in 20 mM Tris-Cl, 100 mM NaCl, pH 7.3.

### Fluorescence labeling

The EsxA proteins with Cys mutations were treated with 20 mM dithiothreitol (DTT) on ice for 20 min. DTT was removed by passing through a Sephadex G-50 column that was pre-equilibrated in 20 mM Tris-Cl, 100 mM NaCl, pH 7.3. The proteins were concentrated to ~2 mg/ml using a 5-kDa-cutoff vivaspin concentrator (Vivascience). The proteins were incubated with 20-fold molar excess of either IANBD [*N*,*N* ′ -dimethyl-*N*-(iodoacetyl)-*N*’-(7-nitrobenz-2-oxa-1,3-diazol) ethylenediamine or Alexa Fluor 546 (AF546) maleimide (Molecular Probes] for 2 h at RT. The free dye was removed by passing through the G-50 Sephadex column. The fractions containing the NBD-labeled protein were pooled, concentrated and stored at −80°C for future use. The labeling efficiency was calculated as [dye]/[protein]% by absorbance spectrophotometry (ε_478_ = 25,000 M^−1^ cm^−1^ for NBD, ε_554_ = 93,000 M^−1^ cm^−1^ for AF546, and ε_280_ = 17,739 M^−1^ cm^−1^ for EsxA). The measured labeling efficiency for all the mutants was ~ 100%.

### Detection of ASB-14 by LC-MS/MS

To establish the standard curve of ASB-14, ASB-14 standard (purchased from Sigma-Aldrich) was dissolved in MilliQ water at concentration of 0.01, 0.1, 1 and 10 mg/ml. The standard solution was run through a quantitative HPLC, followed by MS/MS to confirm the identity. To detect the ASB-14 contamination in the protein samples, the proteins were precipitated by adding 9 volumes of cold acetone and stored in −20°C overnight. The samples were centrifuged at 15,000 × *g* for 30 min to precipitate the proteins, and the supernatant was transferred to a new tube, dried under nitrogen gas and dissolved in 0.1% AcF3. The samples were further desalted by a C18 desalting column and applied to a quantitative HPLC and MS/MS analysis.

### Time-lapse intensity measurements of NBD emission in live cells

WI-26 cells were detached by 0.25% trypsin and re-suspended into an universal buffer (10 mM HEPES, 10 mM sodium acetate, 10 mM MES, 150 mM NaCl, 2 mM CaCl_2_, 11 mM glucose, 50 mg/L bovine serum albumin, pH 7.3). The cells at OD_600_=0.05 (final concentration) were incubated with 100 μg NBD-labeled EsxA on ice for 1 h in a 2 ml total volume. The cells were then washed twice with cold universal buffer and transferred to a cuvette with a stirring bar in the cuvette holder of an ISS K2 fluorometer (ISS, IL). The temperature of the sample chamber was controlled by a circulating water bath. NBD was excited at 488 nm, and emission was recorded at 544 nm. In addition, a long-pass 510-nm filter was placed before the photomultiplier tube to reduce background scatter of the excitation beam. The total time to monitor NBD fluorescence change was 1 h. Relative Fluorescence change was calculated as the fluorescence intensity at the final time point subtracts the fluorescence intensity at 1,000 second when the temperature setting was changed from 4 °C to 37 °C. For the inhibitor experiments, the cells were incubated at 37 °C with 50 μM cytochalasin D or 1 μM bafilomycin A1 for 35 min before the NBD-labeled EsxA (S35C) protein was added to the cells.

### Confocal fluorescence microscopy

WI-26 cells were plated in a Lab-Tek chamber slide (Nalgen Nunc International, IL) at 3 × 10^4^ cells per well and incubated overnight. The slides were then incubated on ice, and 1:1 mixture of the NBD-labeled EsxA and the AlexaFluro 546-labeled EsxA (Molecular Probes, OR) was added. After incubation for 1 h on ice, the slides were transferred to a humidified CO_2_ incubator at 37 °C. After 60 min, the cell samples were fixed for imaging and visualized under a Zeiss confocal microscope. The intensity of NBD was calibrated by the intensity of AF546.

### Bacterial adherence and invasion assays

The adherence and invasion of Mm strains was measured using WI-26, THP-1, RAW264.7 and A549 cells. The cells were planted in 24-well plates (Corning, NY, USA) at 2 × 10^5^ per well and cultured in DMEM media with 10% FBS at 37 °C with 5% CO_2_. After 20 h, the cells were washed twice with PBS and counted. For adherence assay, cells were fixed with 4% paraformaldehyde (PFA) at RT for 10 min and washed with PBS to remove the residual PFA. Then, the fixed cells were infected with single cell bacteria at MOI=1 at 30 °C for 45 min (WI-26, RAW264.7 and A549 cells) or at MOI=5 at 30 °C for 2 h (THP-1 cells). Cells were washed twice with PBS to remove non-adherent Mm. For invasion assay, cells were not fixed with 4% PFA and the infection was the same as adherence assay. After the PBS wash, the DMEM containing 100 μg/ml Amikacin and 2% FBS was added for 2 h to kill extracellular Mm. For both of adherence and invasion, the infected cells were lysed with 200 μl 0.2% (v/v) Triton X-100 (SigmaAldrich, St Louis, MO, USA) solution for 10 min at 30 °C and 100 μl cell suspension was used for ten-fold serial dilutions in PBS. Dilutions (50 μl) were spread on TSA plates and cultured at 30 °C with 5% CO_2_ for 7 days. The CFUs were counted to determine the amount adherent and internalized Mm. Invasion efficiency was calculated as the number of internalized bacteria divided by the number of adherent bacteria.

### Determine the formation of EsxA-ST-SC-GFP complex with fluorescence microscopy and Western blot

Fluorescence microscopy: The indicated Mm strains were cultured in 10 ml 7H9 media with or without Tween-80 and reached to OD_600_ 0.6 ~ 0.8. The bacterial pellet was collected by centrifugation, weighted and resuspended in 1 ml of PBS. 10 μg of the purified SC-GFP was added into each tube and incubated at RT with constant shaking for 1 hr. The bacteria were pelleted by centrifugation and washed with PBS extensively to remove free SC-GFP. Then, the bacterial pellet was resuspended with 1 ml PBS, among which 500 μl of bacterial suspension was loaded to a coverslip and fixed with 4% PFA. The coverslip was immersed in 50% glycerol/water solution and applied to confocal microscopy using a Zeiss LSM 700 (San Diego, CA, USA). Since SC-GFP binds to EsxA-ST with a highly specific covalent bond, the intensity of GFP signal represents the amount of EsxA-ST. To quantify the surface-associated EsxA-ST, the images were analyzed as described before (44). Briefly, the areas with green or red fluorescent signals were first extracted and calculated, respectively. Then, then green areas that are colocalized with red areas were calculated. This green/red ratio was calculated. Layer extraction and area calculation were achieved with Python 3.7.3, according to equation described elsewhere (44,56).

Western blot: The Mm(EsxA-ST) cultured in 7H9 media with/without Tween-80 were collected and incubated with the purified SC-GFP protein at different concentrations (2, 5, 10, 20, 50, 100 μg per 10 mg of bacterial cell pellet) at RT for 1 h in PBS. After extensive washes with PBS, the mycobacterial cells were lysed and applied to Western blot to detect EsxA-ST-SC-GFP by using a mouse anti-GFP antibody (4B10, CST, Danvers, MA, USA) as the primary antibody and the HRP-labelled goat anti-mouse antibody as the secondary antibody. GroEL was detected as the loading control by using an anti-GroEL antibody (CS-44, BEI Resources, Manassas, VA, USA) as the primary antibody and HRP-labelled goat anti-mouse antibody as the secondary antibody. The PVDF membrane was immersed in SuperSignal West Pico (Thermo, Massachusetts, USA), and the chemiluminescent signals were acquired with a iBright FL1500 imaging system (ThermoFisher Scientific, Massachusetts, USA). ImageJ was used to quantify the intensity of chemiluminescence (57). Each group’s relative intensity of EsxA-ST-SC-GFP was calculated according to the intensity from 2 μg SC-GFP per 10 mg of bacterial cell pellet group.

### Calcein AM/Ether-D1 stain and LDH release assays

Calcein AM/Ether-D1 stain assay and LDH release assay were used to determine Mm(EsxA-ST) cytotoxic effect on WI-26 cells. WI-26 cells were planted into a 24-well plate with 1.5 × 10^6^ cells/well and cultured for 24 h. The single cell preparation of Mm(EsxA-ST) cultured with/out 0.05% Tween-80 was used to infect WI-26 cells at MOI=5 for 45 min at 30 °C. After several washes with PBS, the extracellular Mm(EsxA-ST) bacteria were killed by addition of 100 μg/ml Amikacin. The infected cells were cultured at 30 °C for 48 h before the assay.

According to the manufacturer’s manual, the cells were washed and stained with Calcein AM/Ether-D1 solution for 45 min at RT. Then, the excess staining solution was washed, and the cells were imaged in a Floid Cell Imaging Station using the channels for red and green fluorescence, respectively.

The culture supernatant at 48 hours post infection (hpi) was used for LDH release assay. According to manufacture’s manual (CytoTox 96^®^ Non-Radioactive Cytotoxicity Assay, Promega, Madison, WI, USA), 50 μl of cell culture supernatant was mixed with equal volume of CytoTox 96^®^ reagent solution. After incubation at RT for 30 min, another 50 μl of Stop Solution was added to quench the reaction. Absorbance at 492 nm was measured. The cells treated with Lysis Solution were used as the maximum LDH release control. All the acquired data were subtracted with the background reading from the blank cell group. The LDH release percentage was calculated according to equation in the manufacture’s manual.

## Data availability statement

All the date in this study has been included in the manuscript.

## Acknowledgements

The authors appreciate Dr. Hugues Ouellet for providing the pMSP12::mCherry plasmid, and Dr. Jianying Zhang for providing the A549 cells. We thank Dr. Matthias Wilmanns for providing the pMyNT plasmid. The following reagents were obtained through BEI Resources, NIAID, NIH: Polyclonal Anti-*Mycobacterium tuberculosis* Antigen 85 Complex (FbpA/FbpB/FbpC; Genes Rv3804c, Rv1886c, Rv0129c) (antiserum, Rabbit), NR-13800; Monoclonal Anti-*Mycobacterium tuberculosis* GroEL2 (Gene Rv0440), Clone CS-44 (produced *in vitro*), NR-13813.

## Abbreviations

ESX-1: ESAT-6 secretion system-1
EsxA(ESAT-6): 6 kDa early secreted antigenic target
Mtb: Mycobacterium tuberculosis
MPA: Membrane permeabilizing activity
ASB-14: Amidosulfobetaine-14
Mm: Mycobacterium marinum
TB: Tuberculosis
EsxB (CFP-10): 10 kDa cell filtrate protein
HiPPR: High protein and peptide recovery
SDS-PAGE: Sodium dodecyl sulfate polyacrylamide gel electrophoresis
HPLC: High performance liquid chromatography
NBD: C6-Ceramide
WT: Wide type
ATC: Anhydrotetracycline
ST: SpyTag
SC: SpyCatcher
GFP: Green fluorescent protein
LDH: Lactate dehydrogenase
TLR2: Toll-like receptor 2
β2M: β2-microglobulin
ER: Endoplasmic reticulum
PBS: Phosphate saline buffer
CFU: Colony forming unit
GST: Glutathione S-transferase
PCR: Polymerase chain reaction
DTT: Dithiothreitol
AF546: Alexa fluor 546
LC-MS/MS: Liquid chromatography with tandem mass spectrometry
FBS: Fetal bovine serum
PFA: Paraformaldehyde
MOI: Multiplicity of infection
DMEM: Dulbecco’s modified eagle medium
RT: Room temperature
HRP: Horseradish peroxidase
PVDF: Polyvinylidene difluoride
hpi: hours post infection
WB: Westen-blot
CytoD: Cytochalasin D
BA: Bafilomycin A1
DAPI: 4’,6-diamidino-2-phenylindole

